# Accounting for eXentricities: Analysis of the X chromosome in GWAS reveals X-linked genes implicated in autoimmune diseases

**DOI:** 10.1101/009464

**Authors:** Diana Chang, Feng Gao, Andrea Slavney, Li Ma, Yedael Y. Waldman, Aaron J. Sams, Paul Billing-Ross, Aviv Madar, Richard Spritz, Alon Keinan

## Abstract

Many complex human diseases are highly sexually dimorphic, suggesting a potential contribution of the X chromosome to disease risk. However, the X chromosome has been neglected or incorrectly analyzed in most genome-wide association studies (GWAS). We present tailored analytical methods and software that facilitate X-wide association studies (XWAS), which we further applied to reanalyze data from 16 GWAS of different autoimmune and related diseases (AID). We associated several X-linked genes with disease risk, among which (1) *ARHGEF6* is associated with Crohn’s disease and replicated in a study of ulcerative colitis, another inflammatory bowel disease (IBD). Indeed, ARHGEF6 interacts with a gastric bacterium that has been implicated in IBD. (2) *CENPI* is associated with three different AID, which is compelling in light of known associations with AID of autosomal genes encoding centromere proteins, as well as established autosomal evidence of pleiotropy between autoimmune diseases. (3) We replicated a previous association of *FOXP3*, a transcription factor that regulates T-cell development and function, with vitiligo; and (4) we discovered that *C1GALT1C1* exhibits sex-specific effect on disease risk in both IBDs. These and other X-linked genes that we associated with AID tend to be highly expressed in tissues related to immune response, participate in major immune pathways, and display differential gene expression between males and females. Combined, the results demonstrate the importance of the X chromosome in autoimmunity, reveal the potential of extensive XWAS, even based on existing data, and provide the tools and incentive to properly include the X chromosome in future studies.

## INTRODUCTION

Over the past decade, genome-wide association studies (GWAS) have contributed to our understanding of the genetic basis of complex human disease. The role of the X chromosome (X) in such diseases remains largely unknown because the vast majority of GWAS have omitted it from analysis or incorrectly analyzed X-linked data [1]. As a consequence, though X constitutes 5% of the nuclear genome and underlies almost 10% of Mendelian disorders [2–4], it harbors only 15 out of the 2,800 (0.5%) significant associations reported by GWAS of nearly 300 traits [1,5,6]. This 0.5% of associated SNPs is less often in functional loci compared to autosomal associated SNPs [1,5,7], which further suggests that X-linked associations might include a higher proportion of false positives. This is possibly due to most studies analyzing X using tools that were designed for the autosomes [1]. We hypothesize that X explains a portion of “missing heritability” [8,9], especially for the many complex human diseases that exhibit gender disparity in risk, age of onset, or symptoms. In fact, many of the complex human diseases most extensively studied in GWAS are highly sexually dimorphic, including autoimmune diseases [10–12], neurological and psychiatric disorders [13–17], cardiovascular disease [18–22], and cancer [23–26]. Several mechanisms underlying sexual dimorphism have been suggested [12,27–31], including the contribution of the X chromosome [27,32–35]. The hypothesis is further motivated by the importance of X in sexually dimorphic traits in both model organisms and human Mendelian disorders, as well as by its enrichment for sexually antagonistic alleles, which are expected to disproportionately contribute to complex disease risk [36]. Characterizing the role of X in complex diseases can provide insights into etiological differences between males and females, as well as a unique biological perspective on disease etiology because X carries a set of genes with unique functions [37–39].

X-specific considerations that are important to account for in GWAS include, but are not limited to: (1) correlation between X-linked genotype calling error rate and the sex composition of an assay plate, which can lead to plate effects that correlate with sex and, hence, with any sexually dimorphic trait; (2) X-linked variants being more likely to exhibit different effects between males and females [40], suggesting enhanced power of sex-stratified statistical tests; (3) power of the analyses being affected by the smaller allelic sample size (due to males carrying one allele and X-inactivation in females), reduced diversity on X and other unique population genetic patterns [41–47], and a lower density of X-linked SNPs on genotyping arrays; (4) quality control (QC) criteria need to account for sex information to prevent filtering the entirety or a large fraction of the chromosome [1], while at the same time accounting for confounding sex-specific effects; (5) sex-specific population structure leading to differential effects of population stratification (which could lead to false positives [48–50]) between X and the autosomes; and (6) application of association tests designed for the autosomes potentially leading to statistical inaccuracies. Recent advances of association test statistics for X have been made [51–57], with a recent study discovering X-linked loci associated with height and fasting insulin level [56].

Autoimmune diseases (AID) are promising case studies for investigating the role of X in disease because they are commonly sexually dimorphic in symptoms, prevalence (most have higher prevalence in females) [10–12,58], age of onset, and progression [10–12,29,59–62]. While pregnancy [12,30,31] and other environmental factors [63], as well as sex hormones [12,29–31], can contribute to these sexually dimorphic characteristics, a role for X-linked genes has also been suggested [27,62,64–66]. AID have been extensively studied by GWAS, where the majority of autosomal loci discovered have a small effect size, and the combined effect of all associated loci only explains a fraction of heritable variation in disease risk [67–69]. In addition, few of these GWAS have studied the contribution of X and, combined, have provided little evidence for its role in determining disease susceptibility [1,5,6].

In this study, we first introduce X-specific analytical methods and software for carrying out X-wide association studies (XWAS), which take into account several of the above ‘eXentricities’. These methods apply X-specific strategies for QC, imputation, association tests, and tests of sex-specific effects. Furthermore, motivated by the unique characteristics of genes on X, we implemented the first gene-based test for associating X-linked genes and conducted an extensive XWAS of a number of AID and other diseases with a potential autoimmune component [70,71]. Our discovery of X-linked risk genes illustrates the importance of X in AID etiology, shows that X-based analysis can be used to fruitfully mine existing datasets, and provides suitable tools and incentive for conducting such analyses. Additional XWAS can further elucidate the role of sex chromosomes in disease etiology and in the sexual dimorphism of complex diseases, which, in turn, will contribute to improved sex-specific diagnosis and treatment.

## RESULTS AND DISCUSSION

### Datasets and analysis pipeline

We assembled for analysis 16 datasets of AID and other diseases (Table 1). To facilitate independent analysis and replication, we removed individuals from some datasets such that no overlapping data remained between the 16 datasets (Materials and Methods). For each dataset, we first carried out QC that was developed expressly for the X chromosome (Materials and Methods), and excluded the pseudoautosomal regions (PARs). We then imputed SNPs across the X chromosome based on whole-genome and whole-exome haplotype data from the 1000 Genomes Project (Materials and Methods). Of the 16 datasets, none of the original GWAS had imputed variants in an X-specific manner, and only the Wellcome Trust Case Control Consortium 1 (WT1) carried out an analysis of X that is not identical to that of the autosomes [72].

**Table 1.** GWAS datasets. For each of the case-control datasets analyzed in this study, the table lists its name, the disease considered, the number of X-linked SNPs (# *SNPs*), which include imputed SNPs, the number of genes tested in gene-based tests (# *Genes*), and the combined number of SNPs mapped to these genes or to within 15kb of them (# *SNPs in genic regions*). The number of individuals (*# Cases* and *# Controls*) represents the number of samples following QC. The number of males and females in each category is denoted in parenthesis. All datasets consist of individuals of European ancestry. *As control individuals overlap across these datasets, we only considered non-overlapping control subsets for each of the datasets (Materials and Methods). The size of these subsets is indicated under *# Controls.*

| Dataset | Disease | # SNPs | # Genes<br>(#SNPs in<br>genic regions) | # Cases (males,<br>females) | # Controls (males,<br>females) |
| --- | --- | --- | --- | --- | --- |
| ALS Finland [123] | Amyotrophic<br>Lateral Sclerosis<br>(ALS) | 207,947 | 970 (72,219) | 400 (198, 202) | 490 (103, 387) |
| ALS Irish [124] | Amyotrophic<br>Lateral Sclerosis<br>(ALS) | 219,300 | 967 (77,043) | 221 (119, 102) | 210 (112, 98) |
| Psoriasis CASP<br>[128] | Psoriasis | 184,246 | 953 (62,106) | 1,209 (588, 621) | 1,271 (585, 686) |
| Celiac Disease<br>CIDR [125] | Celiac Disease | 187,284 | 962 (64,836) | 1,576 (447, 1129) | 504 (225, 279) |
| CD NIDDK [127] | Crohn's Disease<br>(CD) | 176,072 | 837 (58,874) | 791 (378, 413) | 922 (457, 465) |
| CD WT1* [72] | Crohn's Disease<br>(CD) | 150,275 | 930 (49,017) | 1,592 (607, 985) | 1,701 (923, 778) |
| UC WT2* [131] | Ulcerative Colitis<br>(UC) | 196,781 | 963 (67,422) | 2,341 (1148,<br>1193) | 1,699 (843, 856) |
| MS case control<br>[96] | Multiple Sclerosis<br>(MS) | 183,954 | 842 (61,119) | 943 (312, 631) | 851 (290, 561) |
| MS WT2* [132] | Multiple Sclerosis<br>(MS) | 169,707 | 962 (58,463) | 2,666 (698, 1968) | 1389 (700, 689) |
| Vitiligo GWAS1<br>[126] | Vitiligo | 157,676 | 958 (54,384) | 1,391 (411, 980) | 4,521 (1985, 2536) |
| Vitiligo GWAS2<br>[133] | Vitiligo | 187,688 | 962 (64,974) | 415 (144, 271) | 2,552 (973, 1579) |
| T2D GENEVA<br>[129] | Type-2 Diabetes<br>(T2D) | 220,752 | 971 (75,941) | 2,515 (1050,<br>1465) | 2,850 (1187, 1663) |
| T2D WT1* [72] | Type-2 Diabetes<br>(T2D) | 152,996 | 927 (49,956) | 1,811 (1051, 760) | 1,668 (710, 958) |
| T1D WT1* [72] | Type-1 Diabetes<br>(T1D) | 152,304 | 926 (49,718) | 1,867 (954, 913) | 1,714 (941, 773) |
| RA WT1* [72] | Rheumatoid<br>Arthritis (RA) | 146,907 | 925 (47,880) | 1,772 (443, 1329) | 1,709 (920, 789) |
| AS WT2* [130] | Ankylosing<br>Spondylitis (AS) | 200,042 | 966 (69,441) | 1,472 (976, 496) | 1,260 (665, 595) |

In each of the datasets, we applied three statistical tests for association of each SNP with disease risk: FM_02_, FM_F.comb_, and FM_S.comb_ (Materials and Methods). The FM_02_ test utilizes logistic regression as commonly applied in GWAS, where X-inactivation is accounted for by considering hemizygous males as equivalent to female homozygotes. The other two tests employ regression analyses separately for each sex and combine them into a single test of association using either Fisher’s method (FM_F.comb_) or Stouffer’s method (FM_S.comb_). The FM_F.comb_ test accommodates the possibility of differential effect size and direction between males and females and is not affected by the allele coding in males (i.e. whether each allele in males is counted twice as in FM_02_ or only once; Materials and Methods). FM_S.comb_ takes in account the potentially different sample sizes of males and females and the direction of effect, thereby increasing power in some scenarios (see Supplementary Text). We employed EIGENSOFT [48] to remove individuals of non-European descent and to correct for potential population stratification. Following this correction, QQ (quantile-quantile) plots for each of the three tests across all SNPs, along with genomic inflation factors, revealed no systematic bias across the datasets (Figure S1; Table S1). We provide results for association of individual SNPs with disease risk in Supplementary Text, Figure S2, and Table S2, and focus on the results of gene-based tests (described below) for the remainder of our analysis.

We applied a gene-based test to X-linked genes in each of the 16 datasets using the FM_02_, FM_F.comb_ and FM_S.comb_ statistics. Gene-based tests aggregate association signals across a group of SNPs within a locus while considering the dependence between signals due to linkage disequilibrium (LD) to assign a level of significance for the association of the locus overall. It thereby also reduces the multiple hypothesis-testing burden from the number of SNPs to the number of tested loci [73–76]. This approach can increase power for the autosomes [75,77] and enable replication based on a different set of SNPs in the associated locus. Due to some issues discussed above (see Introduction), this increase in power can be even more pronounced for X.

For our gene-based tests, we defined genes by unique transcripts and included a flanking 15 kilobase (kb) window on each side of the transcribed region to also consider cis-regulatory elements. We used the truncated tail strength [78] and truncated product [79] methods (Materials and Methods) to combine signals across all SNPs in a gene, while accounting for LD. These two methods combine signals from several of the most significant SNPs, thus improving statistical power compared to gene-based tests that consider all SNPs or only the SNP with the strongest signal in the gene. This is especially important for cases in which a gene contains multiple risk alleles or when the causal SNP is partially tagged by multiple tested SNPs [80,81]. From the first round of discovery, we considered for replication genes with a significance of P < 10^-3^ (Tables S3–S4). For these, we first attempted replication in a different dataset of the same disease (including the related Crohn’s disease and ulcerative colitis), if such a dataset was available for analysis (Table 1), and applied Bonferroni correction for the number of genes we attempted to replicate. Otherwise, motivated by the shared pathogenicity of different AID [82–85] (which is also supported by our following results), we attempted replication in all other datasets considered herein (Table 1). In both cases, we attempted replication using the same test statistic that passed the first round of discovery.

### Associations of X-linked genes with autoimmune and other complex diseases

We detected 54 unique genes that passed the initial discovery criterion in one or more of the 16 datasets. Of these, 38 genes were significant based on the FM_02_ test, 22 based on the FM_F.comb_ test, and 34 in the FM_S.comb_ test (Tables S3–S4), with overlap between the three tests due to their statistical dependence. For 42 of these 54 genes, we had an independent dataset for the same or related disease with which to attempt replication. Of these 42 genes, 5 (12%) successfully replicated, with 3 of the 5 both discovered and replicated based on more than one of the three tests (Figure 1a-c and Table 2). These include 3 genes (*FOXP3, PPP1R3F* and *GAGE10*) in LD for the FM_02_ test and 3 genes (*PPP1R3F, GAGE12H* and *GAGE10*) in LD for the FM_S.comb_ test that are associated with vitiligo. To reduce the level of LD, we repeated the gene-based testing without the flanking region of 15 kb around each gene. All genes still successfully replicated in this case, though it remains unclear whether these represent independent signals or remain in LD with the same—likely unobserved—causal variant(s).

**Table 2.** Gene-based associations replicating in similar diseases. All genes with a discovery nominal P < 1x1 0^-3^ (in *Discovery dataset*) that also replicated in a dataset of the same or similar disease (*Replicated dataset*). Results are presented for each of the 3 tests of association, as well as for the test of sex-differential effect size, as indicated by titles in the table. For both discovery and replication, p-values of both methods of gene-based testing (truncated tail strength and truncated product) are presented. Combined p-values (last column) were calculated using Fisher’s method.

| Discovery dataset | Gene | p-value (tail, product) | Replication dataset | p-value (tail, product) | combined p-value (tail, product) |
| --- | --- | --- | --- | --- | --- |
| **FM<sub>02</sub>** |  |  |  |  |  |
| Vitiligo GWAS1 | PPP1R3F | 6.60x10<sup>-5</sup>,<br>1.39x10<sup>-4</sup> | Vitiligo GWAS2 | 8.10x10<sup>-3</sup>,<br>2.70x10<sup>-3</sup> | 8.26x10<sup>-6</sup>,<br>5.93x10<sup>-6</sup> |
| Vitiligo GWAS1 | FOXP3 | 1.11x10<sup>-4</sup>,<br>2.76x10<sup>-4</sup> | Vitiligo GWAS2 | 5.60x10<sup>-3</sup>,<br>5.40x10<sup>-3</sup> | 9.50x10<sup>-6</sup>,<br>2.15x10<sup>-5</sup> |
| Vitiligo GWAS1 | GAGE10 | 1.60x10<sup>-3</sup>,<br>4.03x10<sup>-4</sup> | Vitiligo GWAS2 | 2.80x10<sup>-3</sup>,<br>3.80x10<sup>-3</sup> | 5.97x10<sup>-5</sup>,<br>2.20x10<sup>-5</sup> |
| CD WT1 | ARHGEF6 | 1.70x10<sup>-3</sup>,<br>3.66x10<sup>-4</sup> | UC WT2 | 2.30x10<sup>-3</sup>,<br>3.10x10<sup>-3</sup> | 5.26x10<sup>-5</sup>,<br>1.67x10<sup>-5</sup> |
| **FM<sub>F,comb</sub>** |  |  |  |  |  |
| Vitiligo GWAS1 | PPP1R3F | 1.14x10<sup>-4</sup>,<br>4.96x10<sup>-4</sup> | Vitiligo GWAS2 | 3.70x10<sup>-3</sup>,<br>5.80x10<sup>-3</sup> | 6.61x10<sup>-6</sup>,<br>3.96x10<sup>-5</sup> |
| **FM<sub>S,comb</sub>** |  |  |  |  |  |
| Vitiligo GWAS1 | PPP1R3F | 6.0x10<sup>-6</sup>,<br>7.60x10<sup>-5</sup> | Vitiligo GWAS2 | 4.80x10<sup>-3</sup>,<br>1.30x10<sup>-3</sup> | 5.29x10<sup>-7</sup>,<br>1.69x10<sup>-6</sup> |
| Vitiligo GWAS1 | GAGE12H | 6.34x10<sup>-4</sup>,<br>6.34x10<sup>-4</sup> | Vitiligo GWAS2 | 4.60x10<sup>-3</sup>,<br>4.60x10<sup>-3</sup> | 4.01x10<sup>-5</sup>,<br>4.01x10<sup>-5</sup> |
| Vitiligo GWAS1 | GAGE10 | 1.85x10<sup>-3</sup>,<br>2.66x10<sup>-4</sup> | Vitiligo GWAS2 | 2.90x10<sup>-3</sup>,<br>2.80x10<sup>-3</sup> | 7.05x10<sup>-5</sup>,<br>1.13x10<sup>-5</sup> |
| **Sex Difference** |  |  |  |  |  |
| CD WT1 | C1GALT1C1 | 1.97x10<sup>-3</sup>,<br>2.63x10<sup>-4</sup> | UC WT2 | 1.39x10<sup>-2</sup>,<br>1.14x10<sup>-2</sup> | 3.15x10<sup>-4</sup>,<br>4.11x10<sup>-5</sup> |

**Figure 1.**
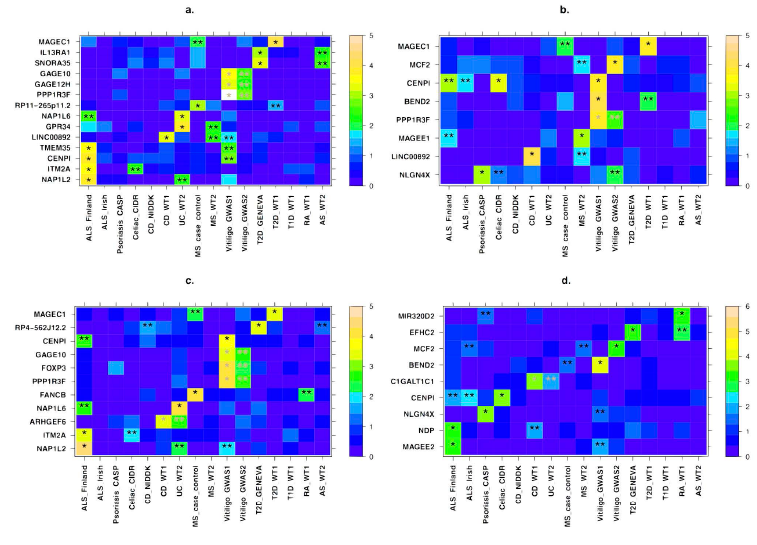
X-linked genes associated with autoimmune disease risk. All genes that showed evidence of association in a gene-based test and replication, including suggestive replication in any other dataset (see main text) are presented for the a) FM_S.comb_ b) FM_F.comb_ c) FM_02_ and d) sex-differentiated effect size tests (Materials and Methods). *X-axis* denotes the different datasets, with their names following the notation from Table 1. *Y-axis* displays the different gene names. For each gene, the more significant p-value of the truncated tail strength and truncated product methods is displayed on a –log10 scale according to the enclosed color scale. A “*” represents the discovery dataset and “**” indicates datasets in which replication is significant after correcting for the number of genes tested for replication. These appear in grey when the discovery and replication are in datasets of the same disease (or across the related Crohn’s disease and ulcerative colitis). Numerical values corresponding to this figure are presented in Tables 2–3.

Of the above four genes we associated to vitiligo risk, *FOXP3* (combined P = 9.5x10^-6^; Table 2) has been previously associated with vitiligo in a candidate gene study of this same dataset [86]. Vitiligo is a common autoimmune disorder that is manifested in patches of depigmented skin due to abnormal destruction of melanocytes. *FOXP3* may be of particular interest as it is involved with leukocyte homeostasis, which includes negative regulation of T-cell-mediated immunity and regulation of leukocyte proliferation [87,88]. Defects in the gene are also a known cause for an X-linked Mendelian autoimmunity-immunodeficiency syndrome (IPEX - immunodysregulation polyendocrinopathy enteropathy X-linked syndrome) [89].

In Crohn's Disease (CD), an inflammatory bowel disorder (IBD) with inflammation in the ileum and some regions of the colon, we discovered association of the gene *ARHGEF6* and further replicated it in the Wellcome Trust Case Control Consortium 2 (WT2) dataset for ulcerative colitis, another IBD (combined P = 1.67x10^-5^). ARHGEF6 binds to a major surface protein of *H. pylori* [90], a gastric bacterium that may play a role in IBD pathology [91,92].

We discovered that another gene, *CENPI*, was associated with three diseases (celiac disease, vitiligo, and amyotrophic lateral sclerosis (ALS)), with an overall combined P = 2.1x10^-7^ (Table S5). The association of *CENPI* remains significant when combining across all 16 datasets and applying a conservative Bonferroni correction for the number of genes we tested (P = 2.7x10^-5^). *CENPI* encodes a member of a protein complex that generates spindle assembly checkpoint signals required for cell progression through mitosis [93]. CENPI is targeted by the immune system in some patients with scleroderma [94]. Additionally, autosomal genes in the same family of genes encoding centromere proteins have been previously associated with ALS (*CENPV*) [95] and with multiple sclerosis (*CENPC1*) [96]. These findings combined suggest a potential pleiotropic role of *CENPI* in risk of AID.

Motivated by the association of *CENPI* in multiple diseases, as well as previous evidence from the autosomes of shared pathogenicity across different AID [82,83], we next sought to replicate the 54 genes from the discovery stage in diseases other than those in which they were discovered. We successfully replicated 17 genes, beyond the aforementioned 5 that replicated in the same or related disease, for a total of 22 (41%) of the 54 genes (Figure 1a-c and Table 3). Six of these 17 were both discovered and replicated based on more than one of the three test statistics, and 5 of the 17 replicated in two separate datasets. We consider these results based on replication in other diseases to provide suggestive evidence of these genes playing a general role in autoimmunity or immune response, and we consider these genes together with the initial 5 in subsequent analyses.

**Table 3.**
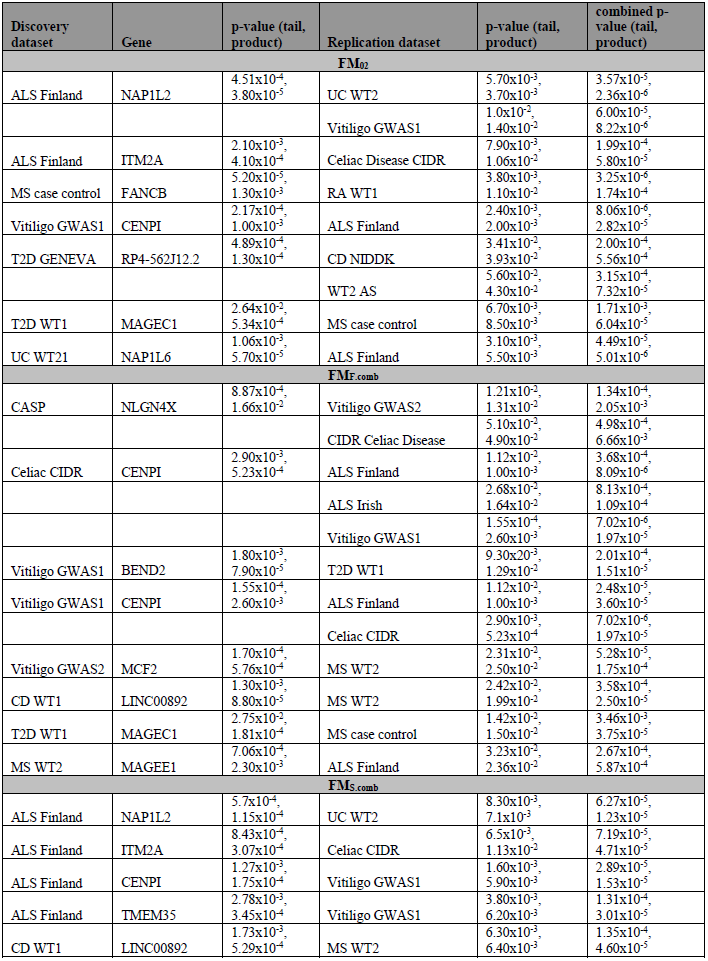
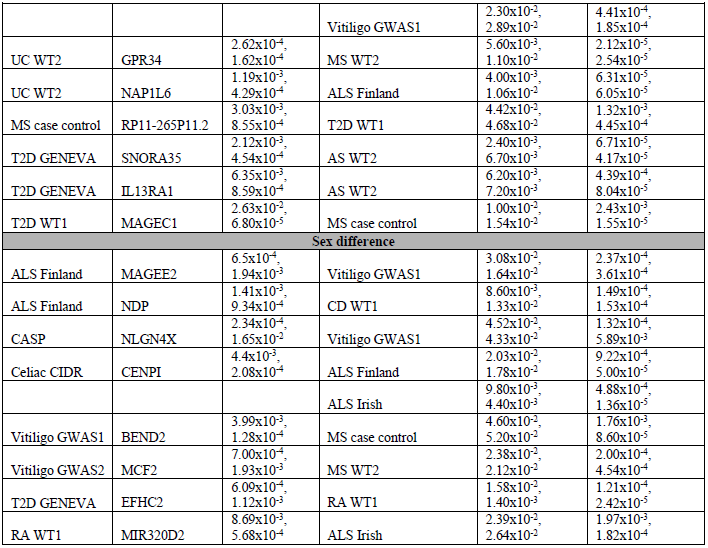
Gene-based associations replicating in other diseases. All genes with a discovery nominal P < 1x10^-3^ that also replicated in a dataset of a *different* disease (see main text). The table mirrors Table 2, with the only difference being whether replication is in the same disease (Table 2) or a different one (this table). Cases in which the same association is replicated in multiple datasets span several rows.

### The sex-specific nature of X-linked genes implicated in autoimmune disease risk

If X-linked genes contribute to sexual dimorphism in complex diseases, then we would expect some genes to have significantly different effect sizes between males and females. We implemented a test of sex-differential effect size (Materials and Methods) and applied it across all SNPs and datasets (Materials and Methods). Consideration of QQ plots and genomic inflation factors revealed no systematic bias (Figure S3; Table S1). As with our above analyses, we combined SNP-level results to a gene-based test of sex-differentiated effect size. This test captures a scenario whereby SNPs within the tested gene display different effects in males and females, without assuming such differential effects to be of a similar nature across SNPs. We followed the same discovery and replication criteria as for the previous analyses, with detailed results provided in Figure 1d, Tables 2–3, and Table S6. Specifically, we discovered and replicated *C1GALT1C1* as exhibiting sex-differentiated effect size in risk of IBD (combined P = 4.11x10^-5^). C1GALT1C1 (also known as Cosmc) is necessary for the synthesis of many Oglycan proteins [97], which are components of several antigens. Defects of C1GALT1C1 may cause Tn Syndrome, a hematological disorder [98]. We also considered replication of sex-differentiated effects in diseases other than the disease of the discovery dataset. This analysis found 8 additional genes, including both *CENPI* (combined P = 1.6x10^-8^) and *MCF2* (combined P = 2.0x10^-4^), which we associated with risk of AID in the above analyses (Tables 2–3). The evidence of sex-differentiated effect of each of these genes is in the same diseases as in the association analysis, thereby pointing to not only a significant contribution of the gene to risk of that disease, but also to its sex-specific effect on the same disease (Figure 1d and Table 3). We again stress that such replication in other diseases is only to be considered as suggestive evidence, and that we consider for subsequent analyses these genes together with those that replicated in the same disease.

Sex-differentiated effects could be a consequence of the X-inactivation (XCI) status of the gene, where at least 25% of human X loci escape XCI to varying degrees. There is no evidence that any of the above three genes (*C1GALT1C1, CENPI*, and *MCF2*) escape XCI [33,99], and all three have degenerate Y gametologs in males; i.e. either the gene has been lost from the Y chromosome (*MCF2*) or the homologous gene on the Y is a non-functional pseudogene (*C1GALT1C1* and *CENPI*). Thus, these genes are expected to show monoallelic expression in both sexes, at least in fibroblasts in which XCI status has been derived [33,99]. Nevertheless, it is possible that these genes show female-biased expression in other tissues as a consequence of escaping XCI in a tissue-specific or disease-specific manner [100,101]. Additionally, the sex-differential risk factor may arise from interaction with other genes and sex-specific environmental factors.

We next directly tested whether any of the X-linked genes that we associated and replicated with AID and related disorders exhibit differences in expression between males and females. We considered a comprehensive dataset of whole blood gene expression from 881 individuals (409 males and 472 females; Materials and Methods) and assayed gene expression in males and females separately. Considering all X genes that we analyzed, they exhibit 2.55-fold enrichment for differential expression between males and females as compared to all genes across all chromosomes (P=6.5x10^-8^). Unsurprisingly, *XIST*, which encodes the long non-coding RNA that induces formation of the Barr body, displays the most significant difference in gene expression between males and females among all X-linked genes (P<<10^-16^). Of the genes we associated and replicated, four exhibit significant sex-differential gene expression: *ITM2A* (4.54x10^-9^), *EFHC2* (4.86x10^-5^), *PPP1R3F* (7.06x10^-5^), and *BEND2* (4.17x10^-4^) (Materials and Methods). Importantly, two of these (*EFHC2* and *BEND2*) also passed initial discovery in the above analysis of sex-differentiated effect sizes, though they were only replicated in datasets different than the one in which they have been discovered (Figure 1d and Table 3). These results suggest that X-linked genes associated with disease risk, especially those that exhibit sex-differentiated effect sizes, are related to sex-differential expression pattern of those genes.

### Association of genes with immune-related function or Y homologs

The nature of the diseases we analyzed and the uniqueness of X led us to an *a priori* hypothesis that genes of a specific biological nature contribute to X-linked AID disease risk. We tested this hypothesis independent of the above results by testing for concurrent association of a whole gene set with each of the individual diseases (Materials and Methods). We tested two different hypotheses by considering 3 such gene sets: The first two sets include X genes with immune-related function as defined by the KEGG/GO or Panther databases (Materials and Methods). The third set includes the 19 non-pseudoautosomal X genes with functional Y homologs. Analysis of the immune-related gene sets was motivated by the nature of the diseases. The test of the last set, on the other hand, was motivated by the evolutionary perspective that genes with functional Y homologs are more likely to be under functional constraint since their Y homologs have survived the progressive degeneration of the Y chromosome over the course of the evolution of the supercohort *Theria* [99]. Thus, they may be more likely to play a part in disease etiology.

The Panther immunity gene set is associated with vitiligo risk in both vitiligo studies that we analyzed and using each of the 3 test statistics of association, as well with type 2 diabetes risk based on the FM_S.comb_ test statistic (Table 4). Similarly, the KEGG/GO set is associated with vitiligo risk in the larger of the vitiligo datasets (Table 4). The set of genes with functional Y homologs suggestively contributes to a much larger group of AID, including psoriasis, vitiligo, celiac disease, Crohn's Disease, and type 1 diabetes, with the first two of these being significant after Bonferroni correction (Table 4). See Table S7 for detailed results for all other datasets and tests.

**Table 4.**
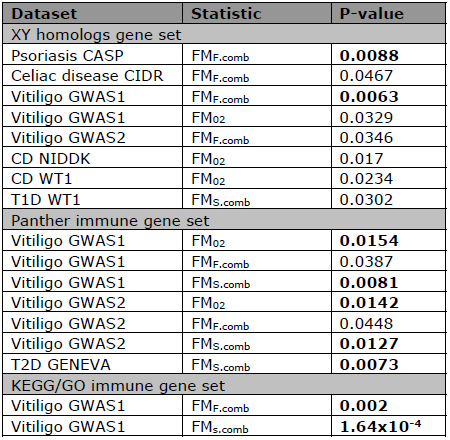
Gene set associations. Three curated gene sets were tested for association with disease risk. Displayed are datasets for which P < 0.05 for association with the gene set indicated in header rows (XY homologs, Panther, KEGG/GO; Materials and Methods). Bold p-values indicate significant associations after multiple testing correction. P-values are the minimum between that based on the truncated tail strength method and the one based on the truncated product method. Results for all datasets and tests are presented in Table S7.

### Relationship and biological functions of genes implicated in autoimmune disease risk

We set out to explore in three analyses the biological function of our associated disease risk genes by considering all 22 protein-coding genes we discovered and replicated with any AID or other complex disease tested. First, we investigated the gene expression patterns of 13 of these genes for which we could obtain tissue-specific expression data (Materials and Methods). Three of these genes show the highest expression in cells and organs directly involved in the immune system (Figures 2-3): *ARHGEF6* is most highly expressed in T-cells, *IL13RA1* in CD14+monocytes, and *ITM2A* in the thymus (in which T-cells develop). Three of the remaining genes, *MCF2* (associated with vitiligo), *NAP1L2* and *TMEM35* (associated with ALS), exhibit the highest expression levels in the pineal gland (Figure 2). The pineal gland produces and secretes melatonin, which interacts with the immune system [102,103] and has been implicated in both vitiligo and ALS [102,104–108], as well as suggested as a possible treatment for ALS [109].

**Figure 2.**
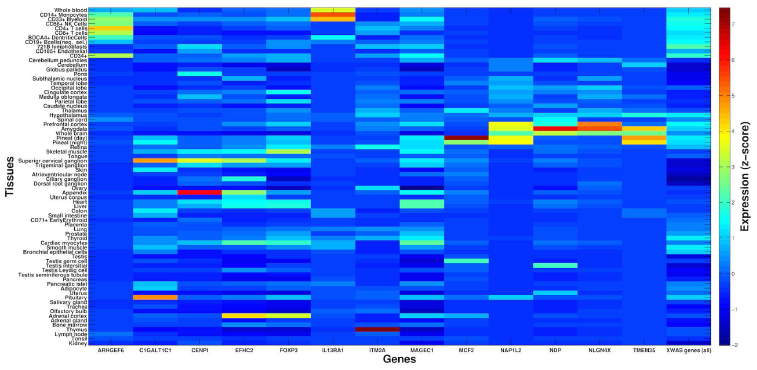
X-linked autoimmune disease risk genes are differentially expressed between tissues. *X-axis* presents 13 of the associated X-linked genes for which gene expression data was available for analysis. For each, a z-score is presented for the deviation of expression in each of 74 tissues (*y-axis*) from the average expression of that gene across all tissues (Materials and Methods). For comparison, the last column shows average z-scores across all 504 X-linked genes that were tested as part of the entire XWAS for which expression data was available. Several associated genes exhibit significantly higher expression in immune-related tissues (see main text and Figure 3).

**Figure 3.**
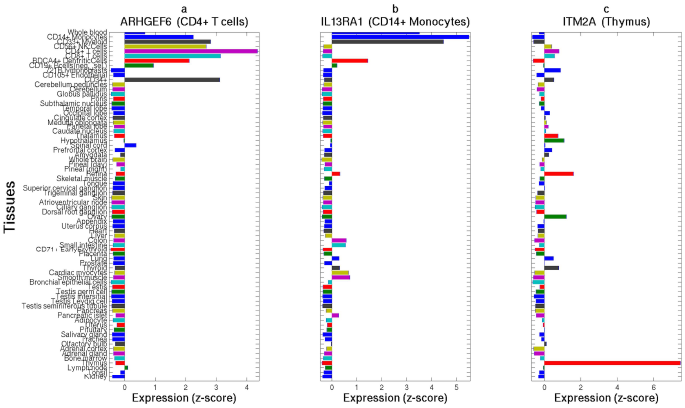
Three X-linked disease risk genes show high expression in immune-related tissues and cells. *ARHGEF6* (a), *IL13RA1* (b), and *ITM2A* (c) show expression greater than 4 standard deviations above the average expression of these genes in T-cells (highest in CD4+ in purple), CD14+ monocytes (blue), and the thymus (red), respectively. *Y-axis* follows the respective tissues from Figure 2 and *x-axis* denotes a z-score for the deviation of expression in each tissue from the average expression of that gene. The title of each panel includes the name of the gene and the tissue with the highest expression for that gene.

Second, we considered co-expression of these 22 associated genes across 881 individuals (Materials and Methods). We observed that 3.9% of all X gene pairs exhibit significantly-positively correlated gene expression patterns. In comparison, 8% of pairs of genes from the set of the above 22 genes exhibit significantly-positively correlated gene expression. This significantly higher fraction relative to X genes overall (Table S8; P=1.53x10^-3^) suggests that genes we associated with disease risk are more likely to work in concert and perhaps interact in the same pathways or cellular networks.

Third, we built an “interactome” by considering this set of 22 protein-coding genes along with genes they interact with in either protein-protein or genetic interactions (Materials and Methods). We found that 18 of these 22 genes are included in the same interaction network (Figure 4), which further supports that they interact with each other. In a pathway enrichment analysis of the resulting interactome (i.e. all genes in Figure 4), several of the significantly enriched pathways relate to immune response or specific immune-related disorders or diseases (Table 5). Another enriched pathway is that underlying lupus, which is a systemic AID. While no dataset for lupus was included in our study, the interactome is potentially enriched for genes in that pathway due to pleiotropy of genes between AID. Other significantly enriched pathways include the regulation of actin cytoskeleton, which can influence the morphology and movement of T-cells, as well as the TGF-beta signaling and ECF-receptor interaction pathways, both of which can mediate apoptosis [110,111]. Finally, the significantly enriched Wnt signaling pathway is generally involved in cell development processes, such as cell-fate determination and cell differentiation [112]. It also plays a role in T-cell and B-cell proliferation and migration, as well as modulation of antigen presenting cells such as dendritic cells [113].

**Figure 4.**
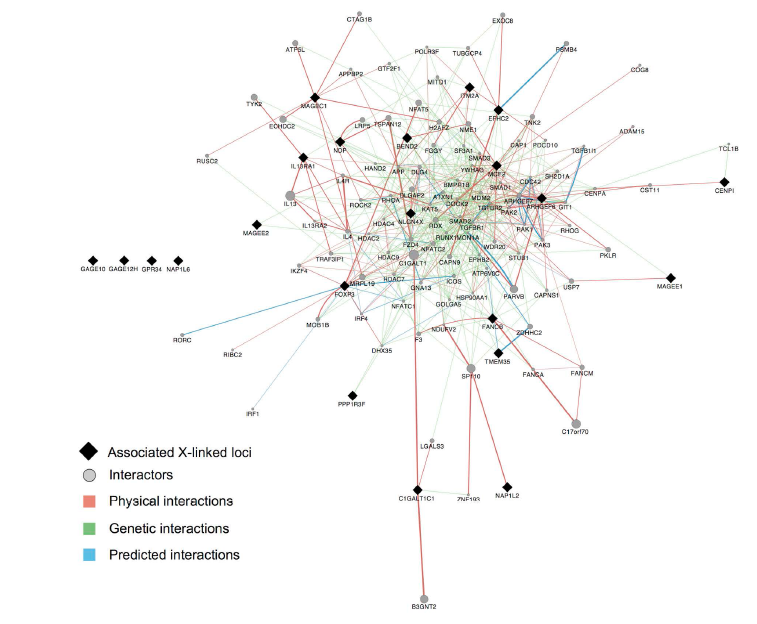
Interactome of X-linked disease risk genes. All 22 X-linked protein-coding genes that showed evidence of association and replication (Figure 1) are denoted by black diamonds and are presented together with genes that interact with them (grey circles) (Materials and Methods). *Physical interactions* refer to documented protein-protein interactions. *Genetic interactions* represent genes where perturbations to one gene affect another. *Predicted interactions* were obtained from orthology to interactions present in other organisms [159]. All but four of these 22 genes share interacting partners according to these known and predicted interactions. Results of a pathway analysis based on this interactome are presented in Table 5.

**Table 5.**
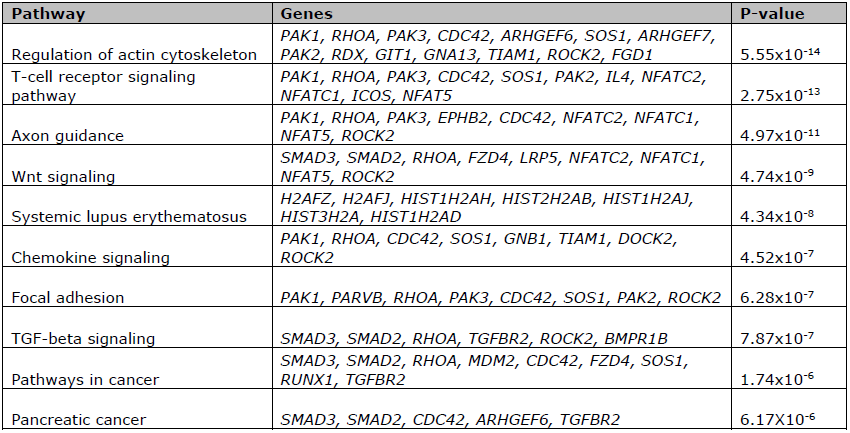
Gene-enrichment analysis of the interactome. Genes we discovered and replicated as associated with any disease tested, and their interacting genes (Figure 4) were enriched for several immune related pathways. We display the ten most significantly enriched pathways. Genes within each pathway that were also within our query set are listed. Displayed p-values are adjusted for multiple testing (Materials and Methods).

### Concluding remarks

In this study, we applied an X-tailored analysis pipeline to 16 different GWAS datasets (Table 1), and thereby discovered and replicated novel associations of several genes with AID risk (Figure 1, Tables 2–3). Multiple lines of evidence point to some of these genes having immune-related functions, including expression in immune-related tissues (Figure 2) and enrichment of these and interacting genes in immune-related pathways (Table 5; Figure 4). Several of the genes we associated with disease are involved in regulation of apoptosis, which plays a role in AID [114–116], including vitiligo [117], psoriasis [118] and rheumatoid arthritis [119]. Our analyses also highlight the sex-specific nature of associated X-linked disease risk genes shedding light on the sexual dimorphism of autoimmune and immune-mediated diseases (Figure 1, Tables 2–3).

The X chromosome has received little attention in the era of GWAS, with growing attention only during the past year [1,56,120,121]. Our results highlight the contribution of X to AID risk and yield new avenues for follow-ups, including unraveling sexual dimorphism in disease etiology. More generally, our study illustrates that with the right tools and methodology, new discoveries regarding the role of X in complex disease and sexual dimorphism can be made, even by mining existing GWAS datasets. Our findings thus underscore the potential for new results and the importance of re-analyzing X in over 2,000 GWAS that have been conducted to date, especially in more recent and better powered studies than the datasets we considered here. To enable such analyses by other researchers, we have made publicly available our X chromosome analysis toolset [122] (http://keinanlab.cb.bscb.cornell.edu) which is in part an extension of PLINK [55].

## Materials and Methods

### Datasets

We obtained 16 GWAS datasets for analysis in this study, which are summarized in Table 1. Datasets were selected to span different autoimmune diseases, including ankylosing spondylitis, celiac disease, Crohn's disease, multiple sclerosis, psoriasis, rheumatoid arthritis, type 1 diabetes, ulcerative colitis, and vitiligo. We also considered datasets of ALS and type 2 diabetes due to suggestive evidence of an autoimmune component to their etiology [70,71].

Out of these, we obtained the following datasets from dbGaP: ALS Finland [123] (phs000344), ALS Irish [124] (phs000127), Celiac disease CIDR [125] (phs000274), MS Case Control [96] (phs000171), Vitiligo GWAS1 [126] (phs000224), CD NIDDK [127] (phs000130), CASP [128] (phs000019), and T2D GENEVA [129] (phs000091).

Additional datasets were obtained from the Wellcome Trust Case Control Consortium (WT): all WT1 [72] datasets, WT2 ankylosing spondylitis (AS) [130], WT2 ulcerative colitis (UC) [131] and WT2 multiple sclerosis (MS) [132] (Table 1). We removed overlapping control samples in order to avoid introducing any biases into replication tests. To accomplish this, we used cases from the WT1 hypertension (HT), bipolar (BP), and cardiovascular disease (CAD) datasets as additional control data. These samples were randomly distributed to the four WT1 datasets, though only BP samples were used as controls for WT1 type 2 diabetes (T2D) due to potential shared disease etiology between T2D, CAD and HT. The WT1 National Birth Registry (NBS) control data was also randomly distributed to the four WT1 datasets. Finally, we randomly distributed the 58 Birth Cohort (58BC) control samples, along with any new NBS samples not present in the WT1 data, between WT2 datasets.

We additionally analyzed the Vitiligo GWAS2 dataset [133], which similar to the Vitiligo GWAS1 dataset that we downloaded from dbGaP, contained case data only. Therefore, we obtained controls from the following datasets in dbGaP: PanScan [134,135] (phs000206), National Institute on Aging Alzheimer’s study [136] (phs000168), CIDR bone fragility [137] (phs000138), COGA [138] (phs000125), and SAGE [138–140] (phs000092). Only samples with the “general research consent” designation in these datasets were used as controls for studying vitiligo. These samples were randomly distributed between the Vitiligo GWAS1 and Vitiligo GWAS2 datasets.

### Quality Control (QC)

Our pipeline for X-wide association studies (XWAS) begins with a number of quality control steps, some of which are specific to the X chromosome. First, we removed samples that we inferred to be related, had > 10% missing genotypes, and those with reported sex that did not match the heterozygosity rates observed on chromosome X [141]. We additionally filtered variants with >10% missingness, variants with a minor allele frequency (MAF) < 0.005, and variants for which missingness was significantly correlated with phenotype (P<1x10^-4^). X-specific QC steps included filtering variants that are not in Hardy-Weinberg equilibrium in females (P<1x10^-4^) or that had significantly different MAF between males and females in control individuals (P<0.05/#SNPs), as well as removal of the pseudoautosomal regions (PARs). We also implemented and considered sex-stratified QC, namely filtering X-linked variants and individuals via separate QC in males and females [120]. However, since we observed no difference in the significant results when applying it to two of the datasets (CD NIDDK, MS case control), we considered data prior to this QC step in our analyses. Finally, following all above QC steps, we removed variants that exhibit differential missingness between males and females (P<10^-7^) [120,142,143]. This step follows the procedure described by König et al. [120] based on a *χ*^2^ test.

### Correction for population stratification

Sex-biased demographic events, including differential historical population structure of males and females have been proposed for many human populations (e.g. [42,46,144–147]). Such sex-biased history is expected to lead to differential population structure on X and the autosomes, thus to differential population stratification. Essentially, population structure on the X captures a 1:2 male to female contribution, while on the autosomes males and females contribute equally to the observed structure. Ideally, population structure on the X needs to be considered to accurately correct for population stratification in an association study of X-linked loci. Hence, we assessed and corrected for potential population stratification via either autosomal-derived or X-derived principal components, and studied the inflation of test statistics in each case as observed in QQ plots. This was performed by principal component analysis (PCA) using EIGENSOFT [48], after pruning for linkage disequilibrium (LD) and removing large LD blocks [50].

For all the datasets analyzed here, which all consist solely of individuals of European ancestry, we found that correction for population stratification is more accurate when based on the autosomes than on X alone due to the smaller number SNPs available to infer structure on X. This observation holds as long as enough autosomal principal components (PCs) are considered. We note, however, that in association studies where more data is available for X, or studies in admixed populations, consideration of population structure on the X chromosome alone can provide a more accurate population stratification correction for XWAS. For example, though African Americans have on average ~80% African and ~20% European ancestry, they exhibit a significant deviation across the X chromosome from these genome-wide estimates. Specifically, African ancestry levels are higher on X, which is due to the sex-biased admixture in which ancestors included relatively more African females and, correspondingly, more European males [148]. Hence, population structure estimated genome-wide (e.g. by EIGENSOFT) may not accurately correct for population stratification in testing X-linked loci in studies of African Americans.

All subsequent analyses are hence based on first excluding any individuals inferred based on EIGENSOFT [48] to be of non-European ancestry. Assessment and correction for population stratification follow the convention of using the first ten autosomal-derived PCs as covariates [49], which is supported by investigation of the resultant QQ plots and by population stratification reported by the original studies. Principal component covariates were not added to the regression model for the ALS Finland, ALS Irish, and CASP datasets as no inflated p-values were observed in these studies [123,124,128] (Figure S1).

### Imputation

Imputation was carried out with IMPUTE2 [149] version 2.2.2 based on 1000 Genomes Project [150] whole-genome and whole-exome (October 2011 release) haplotype data. One of the features added in IMPUTE2 is to account for the reduced effective population size (*Ne*) of the X chromosome by assuming that it is 25% less than that of the autosomes, thereby improving imputation accuracy on the X chromosome. As recommended by the authors IMPUTE2, *Ne* was set to 20,000 and variants with MAF in Europeans < 0.005 were not imputed. Based on the output of IMPUTE2, we excluded variants with an imputation quality < 0.5 and variants that did not pass the above QC criteria (see *Quality Control*). Table 1 displays the number of SNPs we considered in each dataset following imputation and these additional QC steps.

### Single-SNP association analysis

We considered 3 tests for associating X-linked SNPs with disease risk. The first test effectively assumes complete and uniform X-inactivation in females and a similar effect size between males and females. In this test, females are hence considered to have 0, 1, or 2 copies of an allele as in an autosomal analysis. Males are considered to have 0 or 2 copies of the same allele, i.e. male hemizygotes are considered equivalent to female homozygous states. This test is implemented in PLINK [55] as the *–xchr-model 2* option, termed FM_02_ in this study. We do note that the assumptions of complete X-inactivation and equal effect sizes often do not hold (see also our tests and results of sex-differentiated effect size and sex-differentiated gene expression). Hence, in the second test, termed FM_F.comb_, data from each sex (cases and controls) are analyzed separately (with males coded as either having 0 or 2 copies of an allele as above). The female-only and male-only measures of significance are then combined using Fisher’s method [151]. This test accommodates the possibility of differential effect size and direction between males and females and is not affected by the allele coding in males (e.g. 0/2 copies or 0/1 copies). Finally, the third test, termed FM_S.comb_, mirrors the second test except for using a weighted Stouffer’s method [152] instead of Fisher’s method. While Fisher’s method combines the final p-values, Stouffer’s method allows combining and weighing of test statistics. The male-based and female-based test statistics are weighted by the square-root of the male or female sample size [153] and combined while also taking into account the direction of effect in males and females. Implementation follows the equations as provided by Willer *et al.* [153]. Power calculations for these 3 test statistics for a few simulated examples are provided in the Supplementary text.

### Gene-based association analysis

Based on all single-SNP association tests, we implemented an equivalent gene-based test for each statistic by considering all SNPs across each gene. This was carried out in the general framework of VEGAS [73], where the significance of an observed gene-based test statistic is assessed from the distribution of test statistics that is expected given LD between SNPs in that gene [73]. Specifically, the *n* observed SNP-level test statistics are summed together, where *n* represents the number of SNPs in a gene. Next, simulated statistics are obtained as follows: *n* statistics are randomly drawn from a multivariate normal (MVN) distribution and summed. The MVN distribution has a mean of 0 and an *n* x *n* covariance matrix corresponding to the pairwise LD between SNPs mapped to the gene. This procedure is repeated *k* times in order to obtain a distribution of gene-based statistics. The significance is then calculated as the proportion of the *k* simulations that produced statistics that were as or more extreme than the observed one.

Here, we have implemented a slight modification to this procedure: Instead of summing the SNP-based test statistics themselves, we combined SNP-based p-values with either the truncated tail strength [78] or the truncated product [79] method, which have been suggested to be more powerful in some scenarios [80,81]. The simulation procedure is carried out as above, where simulated p-values are derived from the simulated test statistics. To increase time efficiency of the simulation procedure, *k* was determined adaptively as in VEGAS [73].

We obtained a list of X-linked genes and their positions from the UCSC “knownCanonical” transcript ID track (http://genome.ucsc.edu/cgi-bin/hgTrackUi?db=hg19&g=knownGene). SNPs were mapped to a gene if they were within 15 kb of a gene’s start or end positions. When several genes in LD show a significant signal, we repeated analysis while removing the flanking 15 kb on each side.

### Test of sex-differential effect size

In a fourth test, we assayed the difference in the effect size between males and females at each SNP based on statistics derived from the sex-stratified test described above. Considering the female-only and male-only statistics, differential effect size is tested using the following t-statistic [154]:

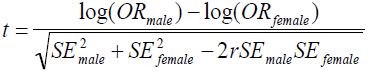

where *OR* stands for the odds ratio estimated in either the male-only or female-only test, *SE* is the standard error in either test, and r the Spearman rank correlation coefficient between log(*ORmale*) and log(*ORfemale*) across all X-linked SNPs. For the odds ratios to be comparable, the odds ratio in males is estimated with coding as having 0 or 2 copies. Finally, we combined the single-SNP tests in each gene into a gene-based test of sex-differential effect size along the same lines as described above for the association test statistics.

### Tests of sex-difference and correlation of gene expression

Whole blood gene expression data for 881 samples (409 males, 472 females) from the Rotterdam Study III [155] was downloaded from Gene Expression Omnibus [156] (accession GSE33828). Expression data was available for 803 of the genes studied in our XWAS. Using a hypergeometric test, we assayed whether the 803 X-linked genes analyzed in our study are more often differentially expressed between males and females as compared to all genes genome-wide. For each gene, we then tested for differential expression between males and females using the Wilcoxon rank sum test across individuals and applied Bonferroni correction to its p-values. We assessed whether any of the 22 protein-coding genes that were associated and replicated in any dataset (Figure 1; Tables 2–3) showed significant sex-differential expression. Expression data is available for 20 of these genes, and Bonferroni correction was applied based on 20 tests.

We tested for co-expression between X-linked genes using the non-parametric Spearman’s rank correlation test between the expression of each pair of genes across the set of 881 individuals. Enrichment of significant co-expression within the set of 20 genes as compared to all 803 genes was tested using a hypergeometric test.

### Tissue-specific gene expression

For analysis of tissue-specific gene expression, we obtained the Human GNF1H tissue-specific expression dataset [157] via the BioGPS website [158]. After excluding fetal and cancer tissues, we were left with expression data across 74 tissues for 504 of the genes studied in our XWAS, including 13 of the 22 genes that were associated and replicated in any dataset. For each gene, we obtained a normalized z-score value for its expression in each tissue by normalizing its expression using the average and standard deviation of the expression of that gene across all tissues.

### Network analysis

A network of interacting genes was assembled in GeneMANIA using confirmed and predicted genetic and protein interactions [159] with a seed list of the 22 protein-coding genes that were associated and replicated across all datasets (Figure 1; Tables 2–3). To minimize bias towards well-studied pathways, all gene-gene, protein-protein and predicted interaction sub-networks were given equal weight when combined into the final composite network. The resulting composite network consisted of the 22 seed genes and the 100 genetic, protein-protein, and predicted interactors with the highest interaction confidence scores. A list of unique genes within this interactome was extracted as input to WebGestalt [160, 161] to discover the ten most significantly enriched pathways in the KEGG database [162]. Enrichment was assessed with the hypergeometric test [160] and reported p-values were adjusted for multiple testing using the Benjamini-Hochberg FDR correction as suggested for such analyses [160]. Pathways that only included a single gene from our interactome were excluded.

### Gene set tests

We additionally tested whether SNPs in a pre-compiled set of genes were collectively associated with disease risk. To accomplish this, we modified the gene-based analysis described above to consider multiple genes. Specifically, the simulation step now entails drawing from *m* different multivariate normal distributions, with *m* denoting the number of genes in the tested gene set. Each of the *m* multivariate normal distributions denotes one gene and has its own covariance matrix that corresponds to the LD between SNPs in that gene. To verify that this procedure, previously proposed for gene-based tests, can be applied to gene sets, we compared p-values derived from phenotypic permutations to this simulation procedure, which revealed highly correlated significance values (Figures S4–S5). Thus, the results we report are based on the simulation procedure, rather than from a limited number of computationally-intensive permutations.

We applied this test to 3 sets of genes: (1) We manually curated a set of immune-related genes from the KEGG [162] pathways and Gene Ontology (GO) [163] biological function categories. We first considered all genes from the two databases in 15 and 14 categories, respectively, that are particularly relevant for autoimmune response. We subsequently removed eight genes from this list that we found were either too general (e.g. cell cycle genes) or too specific (e.g. F8 and F9 blood coagulation genes) to obtain a final list of 27 genes (Table S9); (2) The Panther immune gene set was obtained by including all genes in the category of “immune system processes” in the Panther database [164]; and (3) The XY homolog gene set was obtained from data provided by Wilson-Sayres & Makova [99].

## ACKNOWLEDGEMENTS

Some of the datasets used for the analyses described in this manuscript were obtained through dbGaP accession numbers phs000344, phs000127, phs000274, phs000171, phs000224, phs000130, phs000019, phs000091, phs000206, phs000168, phs000138, phs000125 and phs000092. We thank the NIH data repository, the contributing investigators who contributed the phenotype data and DNA samples from their original study, and the primary funding organizations that supported the contributing studies.

This study also makes use of data generated by the Wellcome Trust Case Control Consortium. A full list of the investigators who contributed to the generation of the data is available from www.wtccc.org.uk.

## SUPPORTING INFORMATION LEGENDS

**Figure S1. QQ-plots for single marker association tests.** Blue triangles denote association p-values for the FM_F.comb_ test, red crosses denote p-values for the FM_S.comb_, while the black points denote association p-values for the FM_02_ test. P-values are plotted on a log scale. Respective genomic inflation factors are summarized in Table S1.

**Figure S2. Significant SNP associations.** (a) A Manhattan plot of the nominal p-values for the FM_02_ (upper), FM_F.comb_ (middle), and FM_S.comb_ (lower) tests of association for chromosome X SNPs in the 16 datasets. The dotted purple lines correspond to the X-chromosome-wide significance threshold for each dataset. The significant associations are shown as red diamonds. (b-c) Regional association plots of the association results of the FM_02_ test and LD for (b) Vitiligo GWAS1 dataset and (c) WT2 UC dataset. LD structure was plotted using a revised version of the *snp.plotter* software [165]. Due to the large number of SNPs in the associated region of Vitiligo GWAS1, only 1 in every 10 of the non-significantly associated SNPs is shown. We focus on regions presented in (b) and (c) since they show the typical LD peaks around significant association signals.

**Figure S3. QQ-plots for test of sex-differentiated effect size.** Similar to Figure S1, except that p-values are for the test of differential effect size between males and females. Respective genomic inflation factors are summarized in Table S1.

**Figure S4. Simulation versus permutation derived p-values for gene-set tests for FM_02_.** Comparison between simulation derived (*x-axis*) and permutation derived (*y-axis*) p-values for the gene-set association analysis using the FM_02_ test statistic. *r* represents Pearson’s correlation coefficient and the significance of the correlation is indicated in parentheses in scientific notation.

**Figure S5. Simulation versus permutation derived p-values for gene-set tests for FM_F.comb_.** Similar to Figure S4 except for considering the FM_F.comb_ test statistic.

**Table S1.** Genomic inflation factors were calculated from the observed p-values in the various tests. No inflation factor exceeds 1.14. Together with the respective QQ-plots (Figures S1 and S3) these results suggest little to no inflation in the observed SNP-level p-values.

**Table S2.** All significant associations (adjusted P < 0.05) as observed in Figure S2. P-values are Bonferroni adjusted for the number of SNPs tested (Table 1).

**Table S3.** All genes with either truncated tail or truncated product P < 1x10^-3^ for the FM_F.comb_ and the FM_S.comb_ tests.

**Table S4.** All genes with either truncated tail or truncated product P < 1x10^-3^ for the FM_02_ test.

**Table S5.** CENPI association p-values for the FM_F.comb_ test across the 16 datasets.

**Table S6.** All genes with either the truncated tail or truncated product P < 1x10^-3^ for the sex difference test.

**Table S7.** All p-values for all gene sets and all datasets are listed. Those with P < 0.05 are highlighted in Table 4 in the main text.

**Table S8.** Pairs of X-linked genes that are significantly co-expressed. Presented are pairs of genes that are significantly co-expressed, after multiple hypothesis correction, along with the squared Spearman’s correlation coefficient (r^2^) and p-value of a Spearman’s rank correlation test (Materials and Methods).

**Table S9.** List of genes in the KEGG/GO immune gene set.

## Supplementary Text

Supplementary information detailing single-SNP association analysis and power calculations for gene-based tests.

